# A molecular dynamics study of antimicrobial peptide translocation across the outer membrane of Gram-negative bacteria

**DOI:** 10.1101/2022.01.20.477041

**Authors:** Pradyumn Sharma, K. Ganapathy Ayappa

**Author notes:** Contributing author.

## Abstract

With rising bacterial resistance, antimicrobial peptides (AMPs) have been widely investigated as potential antibacterial molecules to replace conventional antibiotics. Our understanding of the molecular mechanism for membrane disruption are largely based on AMP interactions with the inner phospholipid bilayers of both Gram-negative and Grampositive bacteria. Mechanisms for AMP translocation across the outer membrane of Gram-negative bacteria composed of lipopolysaccharides and the asymmetric lipid bilayer are incompletely understood. In the current study, we have employed atomistic molecular dynamics and umbrella sampling simulations with an aggregate duration of **~** 8 microseconds to understand the free energy landscape of CM15 peptide within the OM of Gram-negative bacteria, *E. coli*. The peptide has a favourable binding free energy (−130 kJ mol^−1^) in the O-antigen region with a large barrier (150 kJ mol^−1^) at the interface between the anionic core-saccharides and upper bilayer leaflet made up of lipid A molecules. We have analyzed the peptide and membrane properties at each of the 100 ns duration umbrella sampling windows to study variations in the membrane and the peptide structure during the translocation through the OM. Interestingly the peptide is seen to elongate, adopting a membrane perpendicular orientation in the phospholipid region resulting in the formation of a transient water channel during it’s translocation through the bilayer. The presence of the peptide at the lipid A and core-saccharide interface results in a 11% increase in the membrane area with the peptide adopting a predominantly membrane parallel orientation in this cation rich region. Additionally, the lateral displacement of the peptide is significantly reduced in this region, and increases toward the inner phospholipid leaflet and the outer O-antigen regions of the membrane. The peptide is found to be sufficiently hydrated across both the hydrophilic as well as hydrophobic regions of the membrane and remains unstructured without any gain in helical content. Our study unravels the complex free energy landscape for the translocation of the AMP CM15 across the outer membrane of Gram-negative bacteria and we discuss the implications of our findings with the broader question of how AMPs overcome this barrier during antimicrobial activity.

## 1 Introduction

Rising antimicrobial resistance has raised the demand for developing effective antimicrobials. With last-resort antibiotics like polymyxin increasingly showing limited efficacy [1] in clinical settings, there is a pressing need for developing peptide based antimicrobials with a lowered propensity to induce bacterial resistance. The natural origin of certain antimicrobial peptides (AMPs) coupled with a low resistance response in bacteria, renders them as ideal candidates for developing a new class of antimicrobials [2]. Unlike some antimicrobials, cationic AMPs are selective against bacteria due to the inherently anionic nature of the bacterial membrane. AMPs have different mechanisms of disrupting cellular activities and causing cell death. These mechanisms involve cell membrane rupture, hindering cell wall synthesis, inducing osmotic imbalance and compromising the transmembrane potential [3–6].

Antibacterial action of AMPs have been extensively investigated and their mode of entry and organization at the membrane interface is driven largely by electrostatic interactions modulated by the surface charge on the membrane [4, 6]. The balance between charge and hydrophobicity determines their location, orientation and secondary structure conformation within the membrane. At increased peptide to lipid ratios, several mechanisms have been proposed to elucidate the lytic pathways adopted by the peptides. These include pore formation by the barrel-stave and toroidal mechanisms and membrane permeabilization by the carpet mechanism [5, 7]. Although these mechanisms are largely based on the interaction of the AMPs with the inner bacterial membrane, the passage of AMPs through the different barriers in Gram-positive and Gram-negative bacteria is still a matter of debate and relatively unexplored.

The complex bacterial cell envelope, with an outer membrane (OM), periplasmic peptidoglycan (PGN) layer, and cytoplasmic inner membrane (IM) provides a multilayer and multi-barrier environment for antimicrobial translocation [8–13]. The OM illustrated in Fig. 1 is composed of complex lipopolysaccharides (LPS) molecules that provide the initial barrier to antimicrobials molecules [8, 11]. The LPS is topological similar to a polymeric brush having lipid A (LA), acting as an anchor for this macromolecule in the membrane. The LA has hydrophobic tails which face the phospholipid (PL) tails present in the lower leaflet of the OM. The LA is covalently bonded to the core-saccharides (CS), which themselves are further bonded to polymeric O-antigen (OA) sugars. CS and OA are composed of sugar units, and the sugar sequence of these components varies across species and strains [14]. With advances in force-field development and increased computing power, molecular dynamics (MD) simulations have increasingly been used to study the barrier properties of the bacterial cell envelope. Insights into the barrier properties of the bacterial membranes have recently been obtained using free energy methods based on umbrella sampling and metadynamics [8, 10, 11, 15]. The amphiphilic nature of LPS acts as a barrier for both hydrophobic and hydrophilic molecules prohibiting their translocation across the bacterial cell envelope [11]. It has been observed that the OM barrier is asymmetric toward small molecules, and the LPS molecule is found to localize the molecules within this low diffusive environment [8, 11]. While AMPs differ from small molecules, such as benzene and thymol whose free energies of translocation have been studied in the bacterial OM, the translocation and uptake of AMPs are expected to be quite diverse depending on residue, charge and overall size of the peptide. It has been suggested that AMPs affect LPS packing and exhibit a so called “self-promoting” uptake [16, 17], to cross the OM, although the exact mechanism is unclear.

**Fig. 1.**
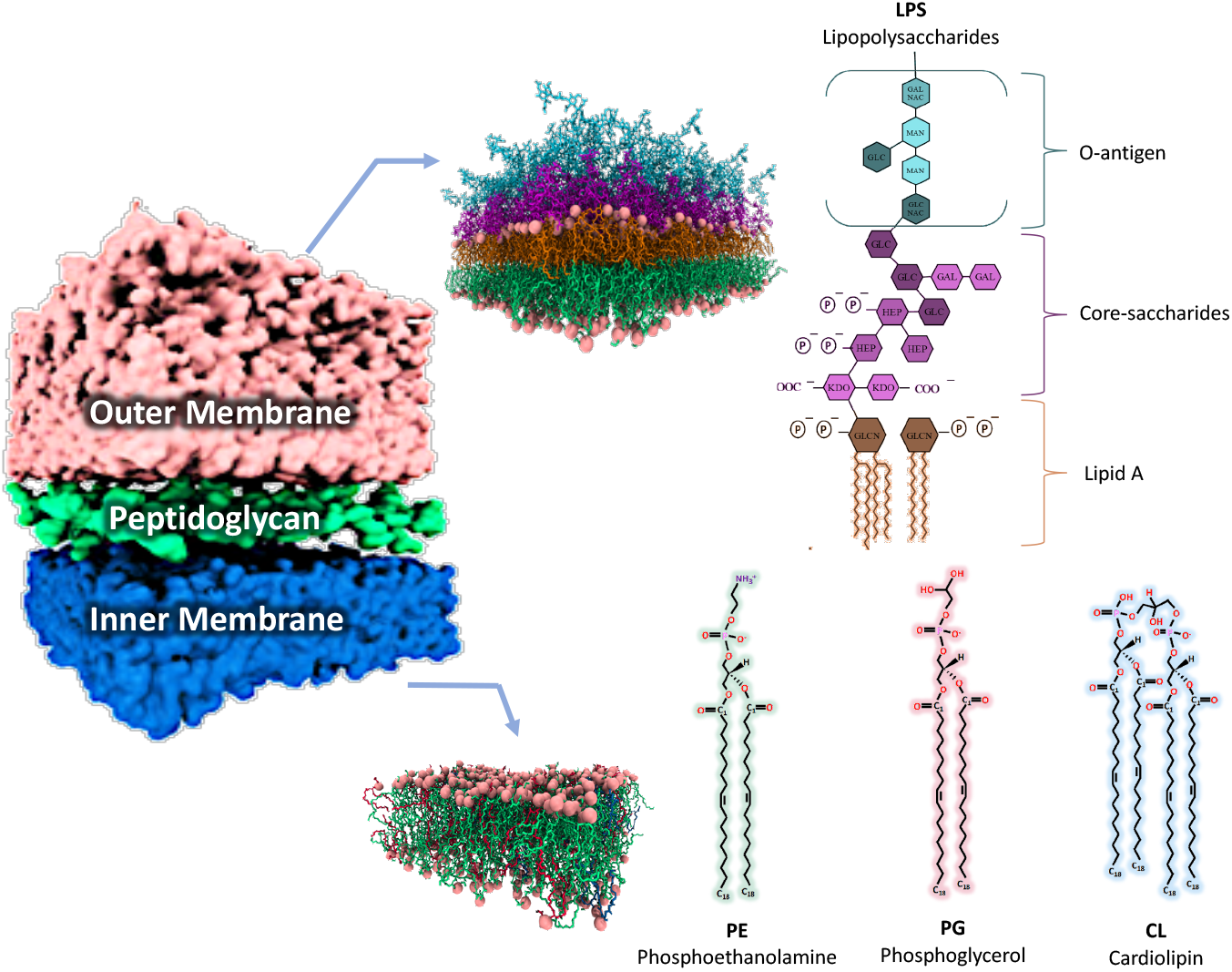
Molecular structures of outer and inner membranes in the cell envelope of Gram-negative bacteria (*E. coli*).

Natural AMPs have their advantages and disadvantages, and hence synthetic AMPs are synthesized to inherit particular traits of the original peptides [18]. Melittin obtained from bee venom has the desired antimicrobial potency but also exhibits hemolytic activity against mammalian cells [19]. Cecropin A, which is obtained from moths, has lowered hemolytic activity against eukaryotic cells [20]. The AMP CM15 (KWKLFKKIGAVLKVL-NH2) is a combination of cecropin A (1-7 residues) and melittin (2-9 residues), to develop an AMP with both antimicrobial potency and minimal hemolytic activity [21]. Widefield fluorescence microscopy has revealed that CM15 induces reactive oxygen species in the cytoplasmic content of bacteria, causing toxicity and exhibiting antibacterial response [22]. Spin labelling electron paramagnetic resonance studies revealed the increased penetration of the N-terminus into the lipid bilayer [23] and sum frequency generation vibrational spectroscopy was used to show that C-terminal amidation increased membrane binding and disruption of DPPG (1,2-Dipalmitoyl-sn-glycero-3-phosphoglycerol) bilayers with CM15 [24].

Molecular dynamics simulations has been extensively used to study binding, translocation and pore forming mechanisms of AMPs through the bilayer phospholipid environment of the bacterial IM. These studies include simulations using both coarse grained [25, 26], atomistic [27–31] or a combination of both methods [32]. Vesicle leakage assays in combination with MD simulations have also been used to determine aggregation states and develop a molecular understanding for the AMP activity [33, 34]. Additionally vesicle based experiments allow the determination of the partitioning coefficients and associated free energies of insertion [35]. In addition to the continued effort to understand AMP activity, there is recent interest in developing antimicrobial polymers as AMP mimics [36]. There have been a few MD studies on the interaction of the CM15 peptide which is the focus of this manuscript, with bilayer membranes. In an all atom study by Bennet et al. [37] the variation of insertion free energies of CM15 into different lipid environments and influence of force-fields has been investigated. Molecular dynamics simulations and free energy computations have been recently used to study the synergy of CM15 uptake in lipid bilayers in the presence of suramin a drug molecule [38, 39] which stabilized the *α*-helical structure of CM15.

With the emphasis on the interaction of AMPs largely restricted to the inner phospholipid bilayer membrane of the bacterial cell envelope, the mechanism of translocation of AMPs through the OM and the PGN layer have not received much attention. Thus the manner in which a peptide molecule passes through the various barriers and arrives at the inner membrane remains an outstanding problem. In this manuscript we turn our attention to translocation of CM15 through the OM of Gram-negative bacteria. Using all atom MD simulations coupled with umbrella sampling free energy computations with an all atom model of the OM, we explore the free energy landscape offered by the smooth (OA inclusive, Fig. 1) LPS incorporated OM model for CM15 translocation. We have calculated the insertion free energy of the CM15 peptide in the OM which includes the passage through the LPS as well as the asymmetric OM. Several structural properties of the membrane and peptide are examined to elucidate the changes induced during the translocation of the peptide through the OM.

## 2 Simulation Details

### MD simulations

The Gromacs 2018.6 MD engine was used to perform molecular dynamics simulations [40]. The CHARMM36 forcefield has been used for modelling the membrane and peptide [41, 42]. Periodic boundary conditions were applied in all three directions. Simulations were performed using a leapfrog integrator with an integration time-step of 2 fs. Nosé-Hoover thermostat [43] (coupling constant of 1.0 ps) and Parrinello-Rahman barostat [44] (coupling constant of 5.0 ps - semi isotropic) were used to perform constant temperature (303.15 K) and constant pressure (1 bar) simulations. The isothermal compressibilities for the barostat were *κ_XY_* = *κ_Z_* = 4.5 × 10^−5^ bar^−1^. Linear constraint solver (LINCS) [45] was used for constraining hydrogen bonds. Particle mesh Ewald (PME) algorithm [46] was employed to compute electrostatic interactions with a cutoff of 1.2 nm. The van der Waals interactions were smoothly switched to zero between 1.0 and 1.2 nm. We have used Visual molecular dynamics 1.9.3 (VMD) for illustrating MD snapshots [47].

### Peptide

The structure of CM15 was obtained from the RCSB PDB database (PDB id - 2JMY) [48]. Upon solvation the helical structure is transformed into a random coil, as observed in previous studies [37]. We have simulated the peptide in an aqueous environment with 3111 TIP3P water molecules [49], eight Na^+^ and 14 Cl^*−*^ ions for maintaining 0.15 M salt concentration. The constant pressure (1 bar - isotropic) and temperature (303.15 K) simulation was performed for 1 *μ*s and the final structure was utilized to perform simulations with the OM.

### Outer membrane

The atomistic smooth LPS OM model was procured from the CHARMM-GUI server [50]. The membrane model represents the *E. coli* membrane having 20 LPS molecules in the upper leaflet, composed of two O111 OA units, R3 core-sugars and LA. The lower leaflet comprises 54 POPE (90%), 3 POPG (5%), and 3 PVCL2 (5%). For electroneutrality of the system Ca^2+^ ions were added in the LPS region, and K^+^ ions were added to the cytosolic side. Additional Na^+^ and Cl^*−*^ ions were added to maintain a salt concentration of 0.15 M. The system has 100 Ca^2+^, 42 Na^+^ and 33 Cl^*−*^ ions. The OM model was hydrated using 13016 TIP3P water molecules [49]. This system was energy minimized and equilibrated following standard six-step CHARMM-GUI protocols [50]. In the final step, the membrane was equilibrated in the NPT ensemble for 20 ns and then further simulated at constant temperature and constant pressure conditions for 200 ns. This membrane configuration was used to perform umbrella sampling simulations. We also performed a restraint-free or equilibrium simulation for a duration of 1 *μ*s with this membrane, by adding 5 CM15 peptides each in the aqueous region above the extracellular leaflet and below the intracellular leaflet of the OM.

### Umbrella Sampling

The CM15 peptide was inserted into the membrane configuration in bulk water in the extracellular region (above the membrane). The free energy of insertion of CM15 peptide in the OM was calculated using umbrella sampling (US) simulations. The collective variable for this calculation is defined as the distance between the center of mass (COM) of the peptide and the COM of the OM along the z-direction. Steered MD simulations were performed to create initial configurations for the umbrella sampling simulations. The peptide was pulled from the bulk water region corresponding to the extracellular environment to the bulk water region in the intracellular environment. The CM15 peptide was pulled at a pulling rate of 0.05 nm/ns to create windows at a spacing of 0.2 nm between 7 nm and −7 nm. We have performed NVT and NPT equilibration at each window for 2 ns and 10 ns, respectively. The run time for each umbrella sampling window was 100 ns, where the peptide was restrained using a harmonic potential with a force constant of 1000 kJ mol^*−*1^ nm^*−*2^. 70 windows were used during the umbrella sampling simulations and adequate overlap of the probability distributions was observed across all the windows. The free energy profile was obtained using the weighted histogram analysis method (WHAM) implemented in Gromacs [51]. The initial 10 ns of umbrella sampling trajectory was discarded from the free energy calculations. However, all other properties are calculated over the entire 100 ns trajectory.

## 3 Results and Discussion

### 3.1 Restraint-free simulation

To understand the interactions between CM15 and the OM of bacteria, we have performed a 1 *μ*s simulation with 10 CM15 peptides. Five peptides were initially placed above and below the the OM (Fig. 2A). During the course of the simulation, we observed the insertion of a few peptides into the lower leaflet of the OM (Figs. 2A and B). The composition of the OM lower leaflet is comparable to the inner membrane of the bacterial cell envelope, and it is composed of phosphatidylethanolamine (PE), phosphatidylglycerol (PG), and cardiolipin (CL) (Fig. 2B). Bennett et al. [37] have observed similar spontaneous penetration of CM15 peptides in phospholipid membranes and found that such a process is stochastic in nature with slow kinetics (relative to currently feasible time scales of all atom MD simulations). We also observed that the peptides initially present above the OM enter the outer OA region, however peptide penetration into the CS region was not observed (Figs. 2A and B). The peptides binding with the inner leaflet and the OA region did not exhibit any secondary structure transitions (Fig. 2A). It has been identified by Bennett et al. [37] that these structural transitions for the CM15 peptide in the membrane environment occurs on the timescale of a few microseconds. Furthermore this transition was observed in a phospholipid membrane which is significantly different when compared with the architecture and chemical nature of the LPS membrane considered here. Over the course of the 1 *μ*s simulation, the density distributions reveal that CM15 predominantly sampled the inner leaflet of the asymmetric lipid bilayer preferring to reside near the lipid headgroups and partially sampled the OA region of the LPS (Fig. 2B). These results reveal the presence of barriers in other parts of the OM for CM15 penetration and translocation. In order to assess the free energy landscape in the OM we performed umbrella sampling simulations.

**Fig. 2.**
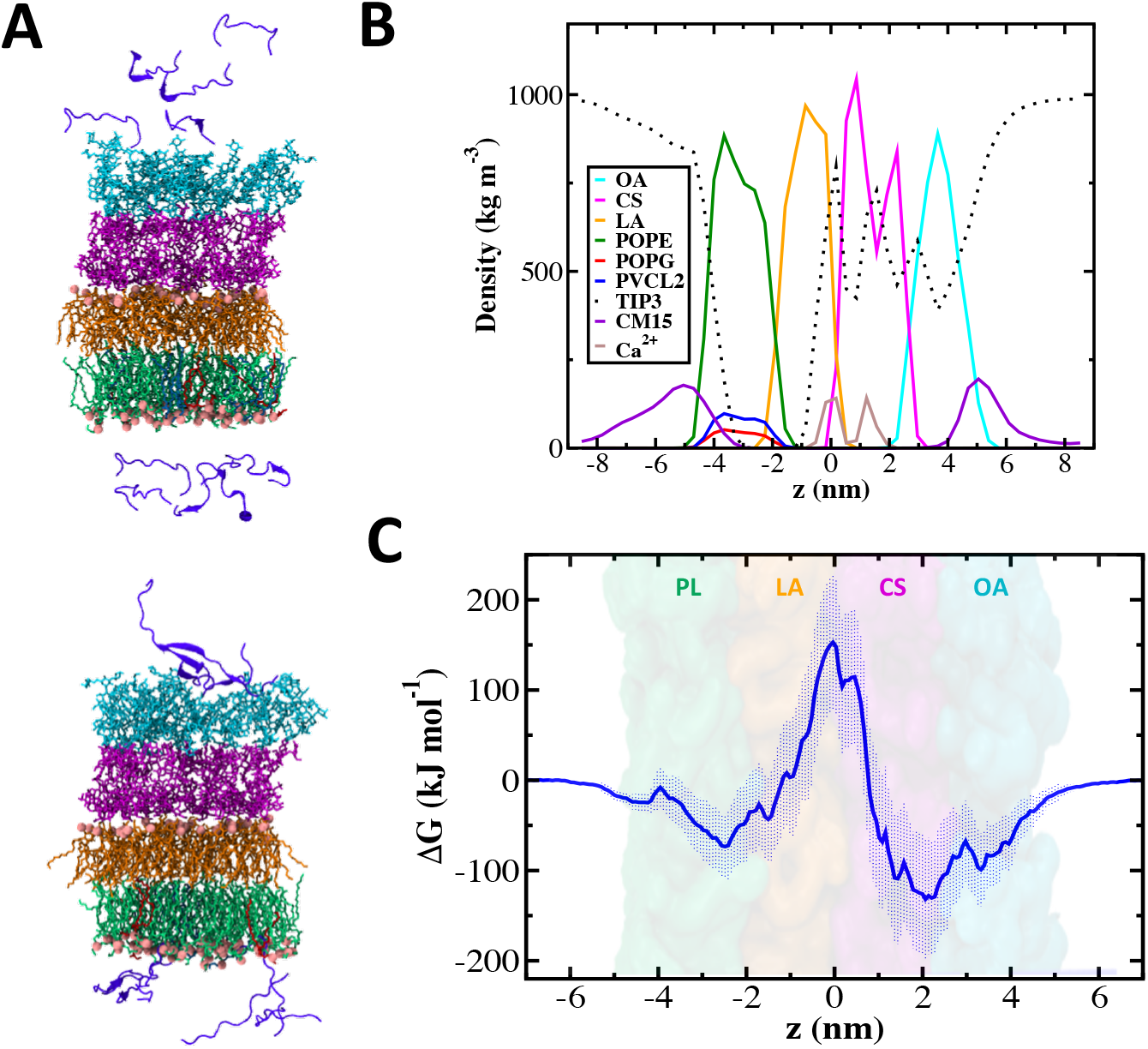
OM offers a significant barrier for the insertion of CM15. (A) Initial (top) and final (bottom) configuration of the 1 *μ*s simulation of the outer membrane and ten CM15 peptides. O-antigen (OA, cyan), core-saccharides (CS, magenta), lipid A (LA, orange), POPE (green), POPG (red), PVCL2 (blue), and CM15 peptides (violet). (B) Density profiles for components of the OM, water, Ca^+2^ ions and peptides along membrane normal z-direction. (C) Insertion free energy of CM15 peptide in the outer membrane of Gram-negative bacteria. Error bars represent the standard deviation. The snapshot (transparent) of the outer membrane is shown in the background with O-antigen (OA, cyan), core-saccharides (CS, magenta), lipid-A (LA, orange), and phospholipids (PL, green). The value of +6 nm and −6 nm corresponds to the extracellular and intracellular environment, respectively.

### 3.2 Free-energy of insertion

To understand the barrier properties offered by the OM of Gram-negative bacteria, we have performed umbrella sampling simulations for the insertion of the CM15 peptide in the OM. The free energy landscape (Fig. 2C) exhibits several interesting features for the peptide translocation process. It is favorable for the peptide to enter the OA region, indicating that peptide binding to the OM is spontaneous as observed in the restraint free MD simulations. The low packing density in the OA region [8] provides a favourable environment for the peptide when compared with the CS-LA interface region [8, 11]. A deep global minima of ~130 kJ mol^−1^ is located close to the OA-CS interface (2-2.5 nm), illustrating that this region offers the most favorable environment for the CM15 peptide. However, a local minima is observed at *z* = 3.3 nm with a barrier of about 25 kJ mol^−1^ for the molecule to move toward the global minima. The presence of this local barrier of about 10 kT is possibly the reason that this region was not fully sampled in the restraint free MD simulations (Fig. 2B).

Though the free energy landscape indicates peptide binding to be a spontaneous process, high energy barriers for the peptide translocation across the OM can also be observed (Fig. 2C). The free energy maxima of ~150 kJ mol^−1^ at the CS-LA interface (*z* = 0 nm) contributes a significant barrier of 280 kJ mol^−1^ for peptide movement into this region from the OA-CS interface (global minima). The presence of this barrier would prevent spontaneous passage of CM15 through the OM at room temperature. The location of this barrier is similar to that observed for the translocation of small molecules such as thymol [8], however the magnitude of the barrier is considerably higher when compared to that of small antibacterial molecules such as thymol where the barrier at the CS-LA interface was 14 kJ mol^−1^. In an all atom simulation of the OM in the absence of the OA, referred to as the rough LPS [11] barriers for hydrophobic molecules such as ethane, benzene and hexane at the LA interface ranged from 12 - 25 kJ mol^−1^. The membrane environment becomes progressively more favourable as the peptide traverses the LA region and enters the PL region of the phospholipid bilayer and a distinct local minima of 75 kJ mol^−1^ is observed in the PL region. The free energy landscape is broad, extending across most of the PL region in the membrane. This situation is unlike the insertion free energy computed for AMPs where the free energy difference between a surface bound folded state and a membrane inserted folded state is positive [35].

In order to understand local orientation of the peptide and the underlying membrane properties we have analyzed various properties of the membrane and the peptide to elucidate the interactions of the CM15 with the OM. We have performed this analysis for all the umbrella sampling windows, which allows us to quantify changes to the membrane properties due to peptide interactions when CM15 passes through the heterogeneous membrane environment.

### 3.3 Membrane Disruption

It is generally believed that the origin of antimicrobial properties of AMPs is due to their membrane disrupting properties [16] with mechanistic interpretations of peptide-lipid interactions restricted to the inner bilayer lipid membrane in bacterial cells. Our umbrella sampling simulations allows us to extract several properties to delineate the influence of the CM15 peptide as it translocates through the OM of the bacterial envelope. To understand how CM15 affects membrane packing, the area per LPS (APL) for each window, defined as the ensemble-averaged lateral simulation box area divided by the number of LPS molecules, was calculated. The APL increases monotonously from the extracellular region to the CS-LA interface (Fig. 3A). The membrane packing is primarily unaffected for the windows where the peptide is present in the OA region and the APL is equivalent to that of the bare membrane system (1.81 +/− 0.02 nm^2^). However, once the peptide enters the CS region, it affects the packing of the membrane, mainly when it is present at the LA and CS interface where the value increases up to 2 nm^2^ inducing a 11% increase in the membrane area. These trends are expected as the OA region is less dense when compared with the tightly packed CS region (Fig. 3A) [8] an unfavourable situation for the CM15 peptide. As the peptide enters the hydrophobic core of the asymmetric bilayer of the OM, changes in the APL of the membrane are reduced, and the APL decreases from the CS-LA interface toward the intracellular region, with larger fluctuations observed in this region. It is unusual to observe an offset between the APL for the umbrella sampling windows corresponding to bulk water in the extracellular and intracellular region. This offset of about 3.3% suggests that extended simulation times during the umbrella sampling might be needed for complete re-equilibration of the membrane area. We have also observed interesting trends when the peptide explores the hydrophobic membrane core region of the OM. The strong interactions of peptide with the headgroup atoms of LA (upper leaflet) and PL (lower leaflet) result in membrane distortions as observed in the snapshots illustrated in Fig. 3C. Irrespective of the proximity of the peptide with the upper and lower headgroup regions, membrane distortions are observed primarily in the lower leaflet (Fig. 3C) concomitant with the formation of a water channel across the bilayer. These findings are similar to previous studies on electroporation of the OM, where the poration defects initiate from the PL region and not the LA region, irrespective of the direction of the electric field [52]. The strong interactions within the LPS molecules in the outer leaflet coupled with the strong hydrophobic interactions due to the six tail structure of the LA, increases the energetic penalty for distortion in the LA headgroups.

**Fig. 3.**
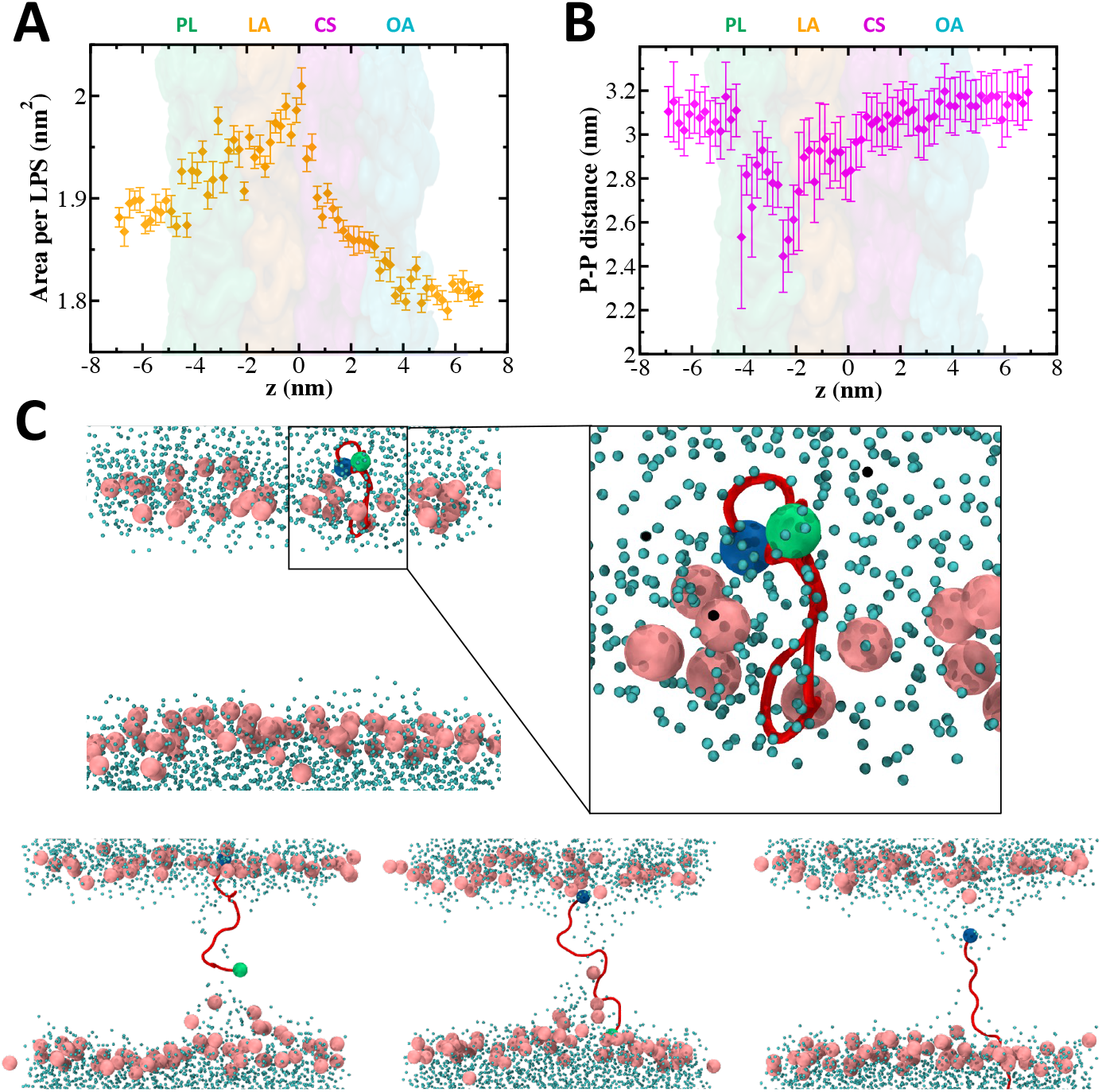
CM15 peptide affects the packing of lipid and LPS molecules in the OM. The variation of (A) area per LPS and (B) minimum interleaflet P-P distance across all umbrella sampling windows. The error bars represent the standard deviations. (C) Configurations showing the distortion in the lower leaflet P atoms for the umbrella sampling windows where the peptide is present in the hydrophobic core of the OM. The other membrane components are hidden for clarity. The peptide, water molecules, and P atoms are shown by red cartoon, cyan beads, and pink van der Waals spheres, respectively. N-terminus and C-terminus of the peptide are highlighted as blue and green beads, respectively. Configurations correspond to umbrella sampling windows at *z* = −0.1 nm (top), −1.3 nm (bottom left), −2.3 nm (bottom center), and −3.7 (bottom right).

To understand these trends in greater detail, we have calculated the minimum interleaflet P-P distance between LA and PL headgroups [53]. The trends show a value of 3-3.2 nm for windows corresponding to OA, CS, and bulk water (Fig. 3B). Interestingly a few windows corresponding to the simulations when the peptide was present in the hydrophobic core of the OM, show up to a 25% reduction (2.4 nm) and a high standard deviation for the minimum P-P interleaflet distance. The reason for these outliers (−4 nm < *z* < 0 nm) is explicitly depicted in a few representative membrane configurations corresponding to these windows (Fig. 3C) where distortions are primarily observed in the PL headgroups. Interestingly, the peptide carries water molecules with it (Fig. 3C), while crossing the hydrophobic region of the OM to create a transmembrane water channel. It is likely that the presence of the 5 Lys residues plays a role in creating the water channel. Since the peptide configurations at the LA-PL interface and regions of the PL represent energetically favourable membrane inserted states (Fig. 2C) the water channel configurations could indeed represent stable configurations of the protein and initiated by the entry of the C-terminus into the membrane from the LA-CS interface. The N-terminus has 4 out of the 5 Lys residues that are present in CM15, belonging to cecropin-A region of CM15 which can drag water present at the LA-CS interface (Fig. 2B) into the hydrophobic region of the membrane. Water channel formation has been observed in coarse grained MD simulations for peptide insertion in bilayer membranes [25, 26] based on the relative orientation of the peptide to the hydrocarbon chains, as well as in early all atom simulations (< 1 ns) of melittin [27] where the protonated N-terminus rich in hydrophilic residues is able to stablize a water channel across the membrane. Interestingly Berneche et al., [27] also reported a loss in helicity in the protonated N-terminus during this relatively short simulation. It is conceivable that thicker transmembrane water channels could be stabilized at high peptide concentrations, resulting in a stable membrane pore. Hitherto we have discussed the alteration in membrane properties due to the presence of the CM15 peptide in different membrane environments and locations sampled in the OM. In the following section, we will discuss peptide reorganization when it traverse through various OM components.

### 3.4 Peptide reorganization

In order to evaluate changes to the peptide structure as it traverses the OM, we have calculated the end-to-end distance, *l_ee_*, and orientation, *θ*, of the peptide as illustrated in Fig. 4. We point out that CM15 remained unstructured and we did not observe any increase in helicity during the entire umbrella sampling simulation. *l_ee_* is defined as the distance between the COM of the N-terminus (LYS-1) and the C-terminus (LEU-15) of the peptide (Fig. 4E). The orientation, *θ* is defined as the angle between the vector joining the center of mass of the N-terminus to the C-terminus with the z-axis of the simulation box. We will discuss these two properties collectively as they provide information on the peptide conformations.

**Fig. 4.**
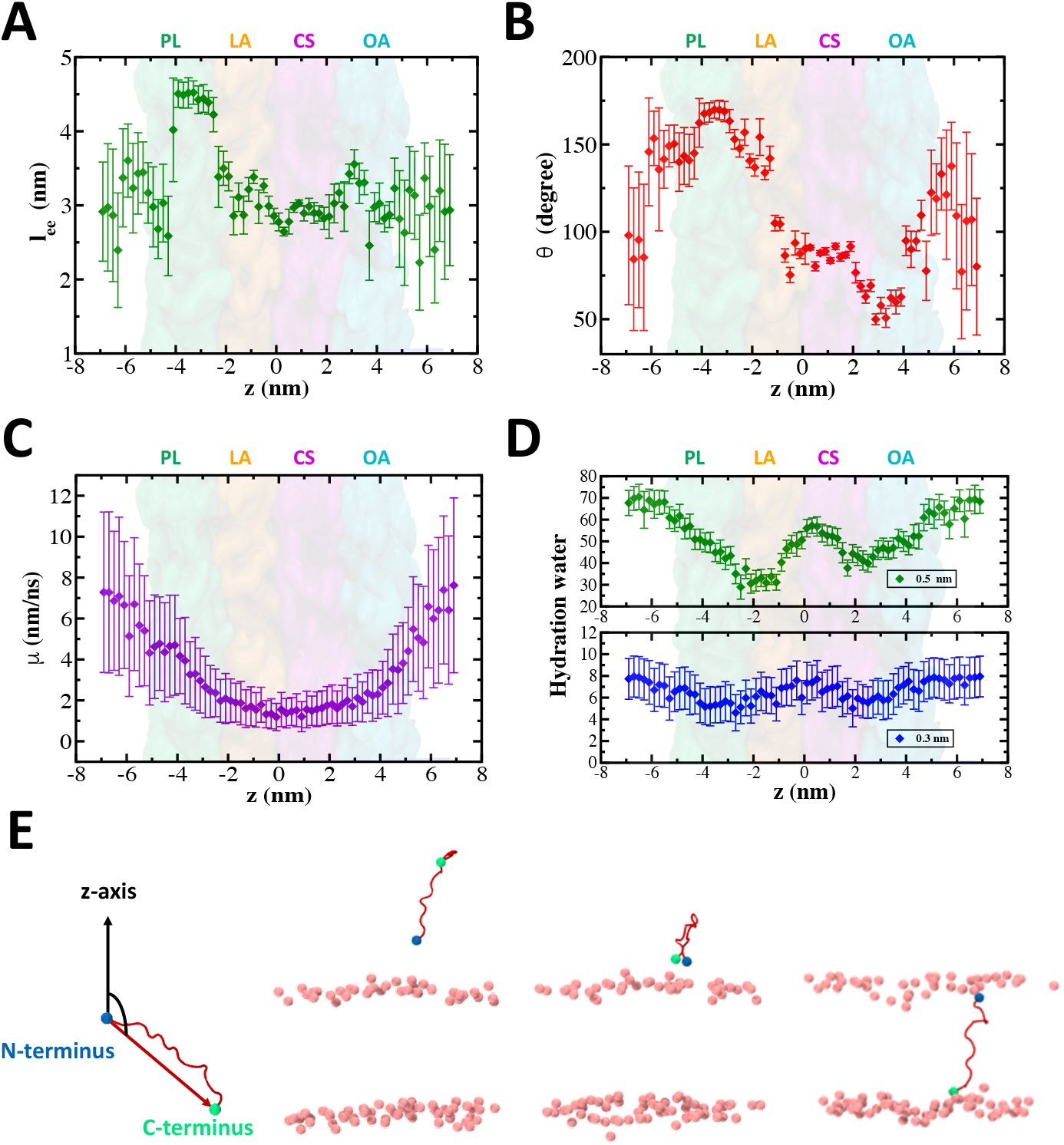
Variation of CM15 peptide internal degrees of freedom and hydration in the low diffusive environment of the OM. The variation of (A) end-end distance, (B) orientation of peptide, (C) lateral mobility, and (D) the number of water molecules present in the hydration shell across umbrella sampling windows. The error bars represent the standard deviations. See the text for detailed definitions of all the collective variables. (E) Schematic defining orientation angle *θ* and membrane configurations highlight the flipping of the peptide. See the text for the definition of the angle. The peptide and P atoms are shown by red cartoon and pink van der Waals spheres, respectively. N-terminus and C-terminus of the peptide are highlighted as blue and green beads, respectively. The configurations correspond to umbrella sampling windows at *z* = 2.9 nm (left), 1.1 nm (center), and −1.9 (right).

In the bulk water environment of both the extracellular and intracellular regions, the peptide effectively sampled a larger configurational space reflected in the high standard deviations for both *l_ee_* and *θ* (Figs. 4A and B). The *l_ee_* of the peptide is ~ 3 nm for all regions except in the lower leaflet PL environment, where an elongated conformation corresponding to *l_ee_* 4.5 nm (Fig. 4A) can be observed. This correlates with the extended configuration formed by the peptide in the lipid region of the membrane where the water channel was observed (Fig. 3C).

The angle *θ* shows greater variation in different components of the OM, varying from ~ 50° in the OA region to ~ 170° in the PL environment. We also observe a flipping of the peptide in the hydrophobic core of the membrane, wherein the orientation of the *l_ee_* vector is reversed as illustrated in (Fig. 4E). This event occurred during the steered MD simulation while the peptide approaches the lipid bilayer from the CS region of the LPS. Since the peptide has reduced rotational degrees of freedom as evident in the low standard deviations for *θ* in the membrane, strong interactions are needed to induce this orientational transition. It appears that the cationic (+2) N-terminus interacts strongly with the anionic LPS molecules, resulting in this change in *θ* as the peptide enters the hydrophobic bilayer region of the OM. Furthermore, this re-orientation enhances favourable hydrophobic interactions between the mostly non-polar residues present toward the C-terminus end of CM15 and the lipids. A value of *θ* = 90° in the CS region illustrates a surface parallel orientation of the peptide in this region. Note that this is the most unfavourable location for the peptide as illustrated in the free energy profile (Fig. 2C). This is in contrast to the other regions where the peptide prefers to adopt configurations that are oriented in a membrane perpendicular configuration. Although the COM of the peptide is located in a region specified by the umbrella sampling window, the components of the peptide are located in neighbouring regions of the OM as well. The extent to which the peptide spreads across the membrane is determined by both *θ* and *l_ee_* and the greatest peptide elongation in the *z* direction is observed in the PL region of the membrane. The peptide orientation plays a significant role during the translocation event due to the heterogeneity of the OM, which provides a hydrophilic environment in OA and CS, and a hydrophobic lipid environment in LA and PL.

In order to evaluate the extent of lateral sampling of the peptide during the umbrella sampling we have calculated the translational x-y “mobility” of the peptide COM to understand the translational degrees of freedom for the peptide. The mobility is defined as [8, 54]

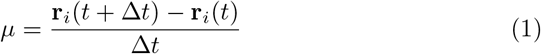

where *μ* is the mobility of peptide and **r**(*t*) is the position in the *x, y* plane at time, *t*. The mobility is a useful metric that allows one to determine the extent of dynamic heterogeneity present in the system. A value of Δ*t* = 1 ns was used in the analysis. Reduced translational freedom is observed in the central regions of the membrane (Fig. 4C). The mobility data shows a symmetric distribution with a minimum located at the CS-LA interface. The mobility of ~7 nm/ns in bulk water drops to a value of ~1.5 nm/ns at the CS-LA interface (*z* = 0 nm). This interface corresponds to the most unfavourable region for the peptide as evident from the free energy barrier at this location (Fig. 2C). The 80% reduction in mobility highlights that the peptide cannot sample the lateral space of the membrane during the umbrella sampling time window of 100 ns reflecting a translationally restricted state of the peptide. We also evaluated the sampling of the lateral x-y region in each of the umbrella sampling windows and found that the sampling was typically within a 5 nm^2^ region around the COM of the peptide. Reduced diffusivity of AMPs have been observed using single molecule particle tracking experiments on supported lipid bilayers with lipid A on the outer leaflet [55]. Measured diffusivities revealed a decrease of about one order when compared with diffusivities observed on symmetric DOPE (1,2-dioleoyl-sn-glycero-3-phosphoethanolamine) supported bilayers. Although we are unable to extract a lateral diffusion coefficient from our simulations, the lowered mobility in the LA region is qualitatively consistent with this observation and we observe a four fold increase in mobility when the peptide is present in the PL region of the bilayer. Both the reduced *l_ee_* and strong electrostatic interactions in the strongly anionic environment of the CS region contribute to the reduced mobility of the peptide. When the peptide adopts a stretched conformation in the bilayer environment (Fig. 4C), the mobility partially recovers indicating that the peptide is able to laterally sample the lipid environment to a greater extent when compared with the CS and LA regions. Similarly when the peptide COM corresponds to the PL region, the stretched conformation (Fig. 4C) illustrates a membrane spanning configuration.

To understand the hydration characteristics of the peptide, we have calculated the number of water molecules within 0.3 nm and 0.5 nm from the peptide atoms using an atom-atom cutoff at each of the umbrella sampling locations (Fig. 4D). It is clear that the presence of hydrophilic Lys groups renders sufficient hydrophilicity to the peptide to hydrate the peptide in all regions of the membrane including the hydrophobic bilayer environment. The variation is more evident with the larger cut-off of 0.5 nm where reduced hydration is observed in the LA environment of the bilayer, with an increase in hydration once the peptide enters the PL region of the lower leaflet. This increased hydration occurs due to the formation of the water channel during the translocation through the bilayer (Fig. 3C). From this analysis it is clear that the peptide has sufficient hydrophilic interactions to carry water molecules even in the hydrophobic core of the membrane (Fig. 4D) leading to the formation of a water channel [25–27]. At the lower cut-off of 0.3 nm there are 6-8 water molecules present in the peptide hydration shell, remaining more or less uniform throughout the membrane environment (Figs. 4D). The presence of this hydrated environment could partly explain the loss of helicity observed for CM15 in the OM.

Before concluding this section we briefly comment on some of the inherent technical issues present while carrying out free energy computations in complex environments such as the outer membrane of Gram-negative bacteria. We have used a single collective variable i.e., the *z* co-ordinate of the center-of-mass of the peptide normal to the membrane bilayer. However a complete exploration of phase space for molecules such as the AMPs might require collective variables judiciously chosen to balance computational feasibility and reliability for studying peptide translocation as discussed recently by Kabekla et al. [26]. The choice of collective variables would also depend on the whether coarse grained force-field [25, 26] are used or whether fully atomistic models [28] are used. Despite these variations we feel that our free energy computations are sufficiently reliable to provide several novel insights into the complex free energy landscape for peptide translocation through the bacterial OM. In order to provide additional support to our findings we carried out a second steered MD pulling simulation for the CM15 through the membrane following the same direction starting from the extracellular region. However we did not carry out an umbrella sampling study for this data set. In this second pulling simulation we observed similar *θ* variations for the peptide as it traverses through the membrane and the flipping event observed at the LA-CS interface was reproducible. However the water channel formation was accompanied by a distortion that occurred primarily in the upper leaflet of LA lipids, in contrast to distortions observed in the lower leaflet in the first pulling simulation. We conjecture that the corresponding free energy for the water channel formed due to distortion in the LA region is likely to result in a greater energetic penalty since the entire LPS molecule along with the CS and OA regions would undergo a distortion along with the cations present in the CS region. The corresponding umbrella sampling simulation would then result in a less favourable free energy change in the lipid environment then that obtained in Fig. 2C which corresponds to the water channel formation driven by lipid displacements in the lower leaflet.

## 4 Summary and Conclusions

To improve the efficacy and selectivity of AMPs, it is crucial to understand their interactions with different components of the bacterial cell envelope. In recent years several molecular dynamics simulations from our laboratory and others have shown that it is important to assess the barrier properties of PGN and OM which are physicochemically and mechanically distinct from the interaction with the bilayer environment of the IM [8–12]. Notably, the OM offers a heterogeneous barrier for antimicrobials. In this study, we have carried out an aggregate of ~8 *μ*s of atomistic MD simulations to understand the interactions of CM15 AMP with the OM of bacteria. The free energy data obtained from umbrella sampling simulations for the translocation of the AMP shows a highly favorable region for the peptide binding at the OA-CS interface. This energy minimum confers a preferable binding site for the peptide. However, the translocation process across the membrane is hindered by a significant barrier (114 kT at room temperature) at the CS-LA interface. Translocation of cationic peptides has been [16, 17] suggested to occur by the displacement of divalent ions present at the anionic binding sites of the CS region of the LPS. A concerted influence of several peptides can potentially lower the barrier for translocation across the CS region giving rise to the so-called “self-promoting” pathway hypothesis. However in our simulations with one CM15 peptide with a net charge of +6, we did not observe this effect and several peptides might be required to understand the charge neutralization effect. Since ions generally take longer to equilibrate we cannot rule out the possibility that extended simulations might be required to observe changes in ion distributions which showed little variation during our umbrella sampling simulations. We also note that the bacterial membrane have several transmembrane protein channels including outer membrane porins, efflux pumps and other transporters whose lipid-protein interfaces could possibly provide lower energy modes for peptide translocation.

Our umbrella sampling simulations provided extensive insight for understanding the alteration in membrane and AMP properties and interaction between these two components when the peptide is present in different environments of the OM. The CM15 peptide alters the packing between the LPS and lipid molecules in the upper and lower leaflets as reflected in the area per LPS computations. The areal changes are the greatest when the peptide is present at the CS-LA interface, showing a distinct maximum corresponding to the maximum in the free energy in this region. We also observed disruption of the membrane headgroups selectively for the lower leaflet phosphorus atoms, irrespective of the proximity of peptide with respect to upper or lower leaflet. This disruption was accompanied by the formation of a transmembrane water channel during the umbrella sampling simulations which correspond to an energetically favorable secondary free energy minima in the PL region of the bilayer. Our simulations suggest that if sufficient charge neutralization with cationic peptides occur to overcome the barrier in the anionic CS region of the LPS the peptide can participate in an energetically favourable process of pore formation. With an increase in the peptide to lipid ratio once can conceive of a synergistic effect wherein peptides cross the LPS barrier and participate in pore formation to form thicker transmembrane water channels in the bilayer, suggestive of a new pathway for AMP action and OM disruption in Gramnegative bacteria. Additionally, the peptide is found to maintain its hydration shell by retaining water molecules in all the configurations sampled in the OM, including those in which the peptide is present within the hydrophobic membrane core.

Our study of CM15 penetration in the OM of Gram-negative bacteria opens up for the first time the complex nature of the interactions between the AMP and various components of the OM. We have studied the interaction of a single peptide with the OM. However, it will be intriguing to observe the synergistic effect of increased number of peptides in future studies. Our study illustrates that it is possible to explore the full complexity of the bacterial cell envelope which includes the outer membrane, peptidoglycan as well as the inner membrane for improved design of antimicrobial molecules.

## Acknowledgments

We thank the Supercomputer Education and Research Center (SERC) for availing computational facility at the Indian Institute of Science, Bangalore. We thank the Department of Science and Technology (DST) and Department of Electronics and Information Technology (DeitY), India for providing funding under National Supercomputing Mission (NSM) and Unilever Research and Development (Bangalore, India) for computing support. Authors would like to thank Prof. Rahul Roy and Satyaghosh Maurya for useful discussions.

## Declarations

Conflict of interest/Competing interests - Authors declare no conflict of interests.

## Notes

### Competing Interest Statement

The authors have declared no competing interest.

